# The full model of the pMHC-TCR-CD3 complex: a structural and kinetics characterization

**DOI:** 10.1101/2020.11.26.397687

**Authors:** Josephine Alba, Oreste Acuto, Marco D’Abramo

## Abstract

The machinery involved in cytotoxic T-cell activation requires three main characters such as: the major histocompatibility complex class I (MHC I) bound to the peptide (p), the T-cell receptor (TCR), and the CD3-complex which is a multidimer interfaced with the intracellular side. The pMHC:TCR interaction has been largely studied both in experimental and computational models, giving a contribution in understanding the complexity of the TCR triggering process. Nevertheless, a detailed study of the structural and dynamical characterization of the full complex (pMHC:TCR:CD3-complex) is still missing, due to insufficient data available on the CD3-chains arrangement around the TCR. The recent determination of the TCR:CD3-complex structure by means of Cryo-EM technique has given a chance to build the entire proteins system essential in the activation of T-cell, and thus in the adaptive immune response. Here, we present the first full model of the pMHC interacting with the TCR:CD3-complex, built in a lipid environment. To describe the conformational behaviour associated with the unbound and the bound states, all atoms Molecular Dynamics simulations were performed for the TCR:CD3-complex and for two pMHC:TCR:CD3-complex systems, bound to two different peptides. Our data point out that a conformational change affecting the TCR Constant *β* (C*β*) region occurs after the binding to the pMHC, revealing a key role of such a region in the propagation of the signal. Moreover, we found that the TCR reduces the flexibility of the MHC I binding groove, confirming our previous results.

## Introduction

T cells play a crucial role in the adaptive immune response. In fact, the TCR triggering is strictly connected to the T cell activation and it is regulated by two main interactions, such as: i) the interaction with the peptide and the MHC class I with the TCR, in the extracellular surroundings; ii) the interaction of the CD3 chains with the TCR in the intracellular environment. The ability of the TCR to recognize a huge number of peptides presented by the MHC ensures a solid defense system from foreign pathogens. The MHC class I is a protein complex formed by an *α* chain, which hosts the binding site for a nine/ten residue-long peptide, and a non-covalent bound *β*2 microglobulin. The pMHC interacts with TCR, a heterodimer composed of two transmembrane glycoprotein chains, named *α* and *β*. The extracellular portion of each chain consists of two domains, the variable region (V) and the constant region (C). The Vα and the V*β* are formed by three complementarity-determining regions, the CDR-loops, which are directly involved in the interaction. The V*α* CDR1 and CDR2 loops are in close contact with the helices of the pMHC complex while the V*β* CDR1 and CDR2 loops interact with the pMHC at the carboxy terminus of the bound peptide. The binding of the TCR to the pMHC implies induced fit conformational changes especially within the V*α* CDR3 loop [1,2]. In addition to the conformational changes in CDRs loops of the TCR, the comparisons between the unliganded and bound forms of TCR-*β*-chain dynamics through NMR spectroscopy showed conformational changes affecting the C*β* regions [3]. Moreover, fluorescence-based experiments and structural analyses have shown that MHC-restricted antigen recognition by the TCR *αβ*-chains results in a specific conformational change confined in the C*α* domain [4]. Therefore, these studies support the view that the remarkable distal changes observed in the C*α* and C*β* regions arise from an allosteric communication mechanism. Such a conformational change is thought to connect the pMHC-TCR information to the CD3 chains, allowing the CD3 ITAM exposure [5,6]. CD3 is a co-receptor protein complex of the TCR formed by four distinct chains organized as dimers: the heterodimers CD3*γ*-CD3*ε*, CD3*δ*-CD3*ε* and the CD3*ζ*-CD3*ζ* homodimer. A previous work [7] has reported a possible stoichiometry of the TCR-CD3 chains of 1:1:1:1, comprising the dimeric subunits such as TCR*αβ*:CD3*γε*:CD3*δε*:CD3*ζζ*. Each subunit of CD3*γ*, CD3*δ*, CD3*ε* present a single extracellular immunoglobulin domain, a membrane-proximal connecting peptide and an intracellular ITAM. Unlike the other chain, the CD3*ζ* has a short extracellular sequence and three ITAM units.

Previous studies have suggested that the interaction of CD3 and TCR *αβ*-chains propagates the TCR–pMHC-binding information to the ITAM regions [5,6] but, although several models of the TCR–CD3 triggering have been proposed, such a mechanism is still largely unknown.

Despite insufficient reliable data on the arrangement of the CD3 chains around the TCR, the determination of the first structure of the TCR interacting with the CD3 chains - recently provided by cryo-EM [8] - has given a chance to build the full construct of the CD3-TCR complex for an in-silico investigation. Therefore, we present the first computational model of the pMHC-TCR-CD3 in a lipid-membrane (Figure 1), providing a complete structural-dynamical description of two different complexes, which differ from the type of peptide bound to the MHC. Of note, we were also able to provide an estimate of the dissociation process between pMHC-TCR interaction in presence of the CD3 construct.

**Figure 1.**
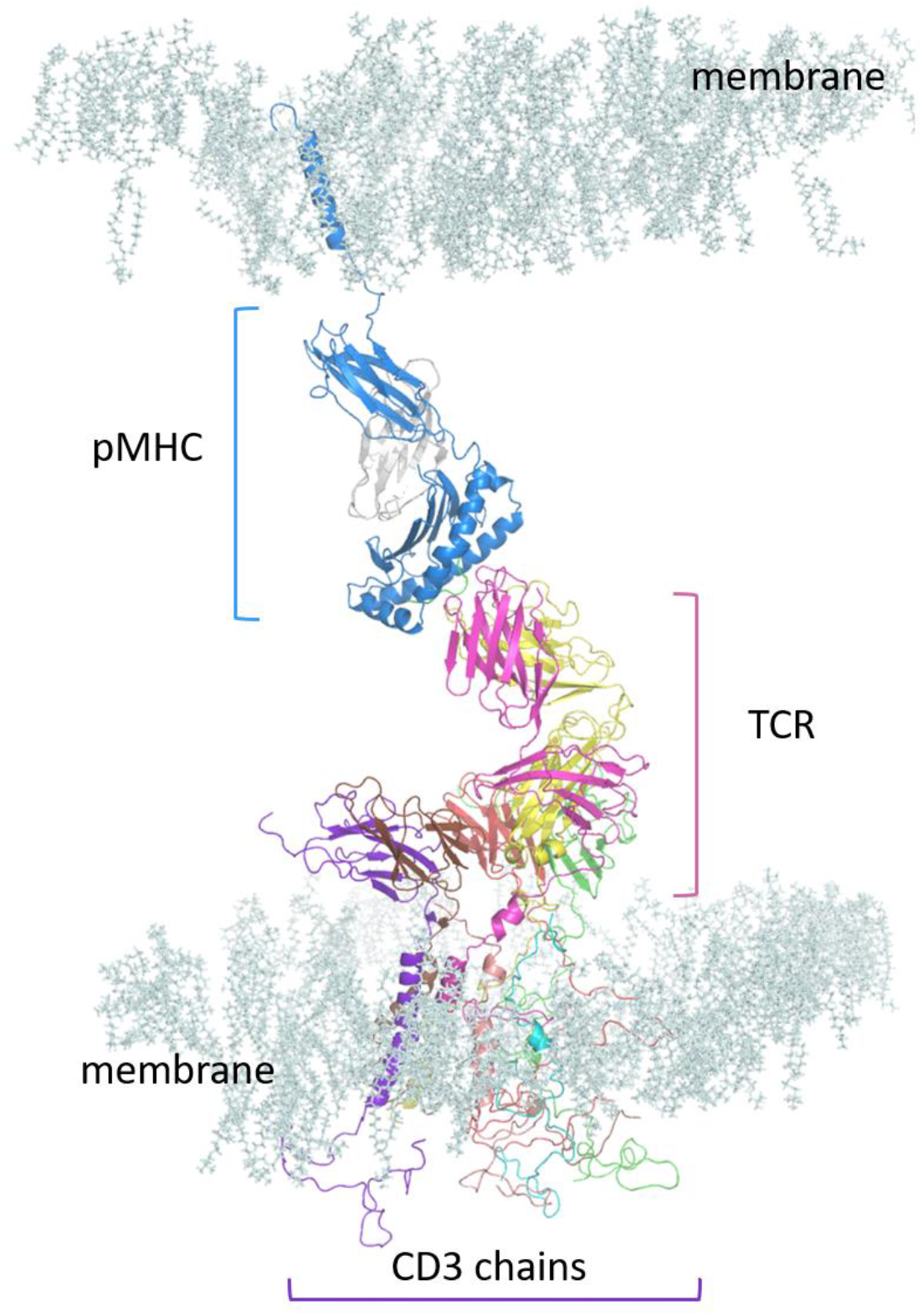
Model of the pMHC:TCR:CD3-chains in membrane. The single regions are labelled. Some lipids were removed in the picture.

## Methods

### Complex modelling

From the cryo-EM structure of the TCR-CD3 complex (PDB ID: 6JXR, resolution of 3.7 Å) the missing parts of the CD3*γε*:CD3*δε*:CD3*ζζ* chains, where built by means of Modeller Software [9], and the final conformations were selected in order to avoid steric clashes (Figure 2). To model the complex with 1G4 TCR, the *αβ* chains provided by the crystallographic construct of such a TCR (PDB ID:2BNR) were aligned to the cryo-EM structure. The transmembrane helices were previously built by means of the Modeller Software [9,10], following the amino acid sequences as provided by the Uniprot database (Uniprot entries P01848 and P01850 for the TCR *α* and *β* chains respectively; Q9MY51 for the *α* chain of the HLA-A* 02:01 [11]).

**Figure 2.**
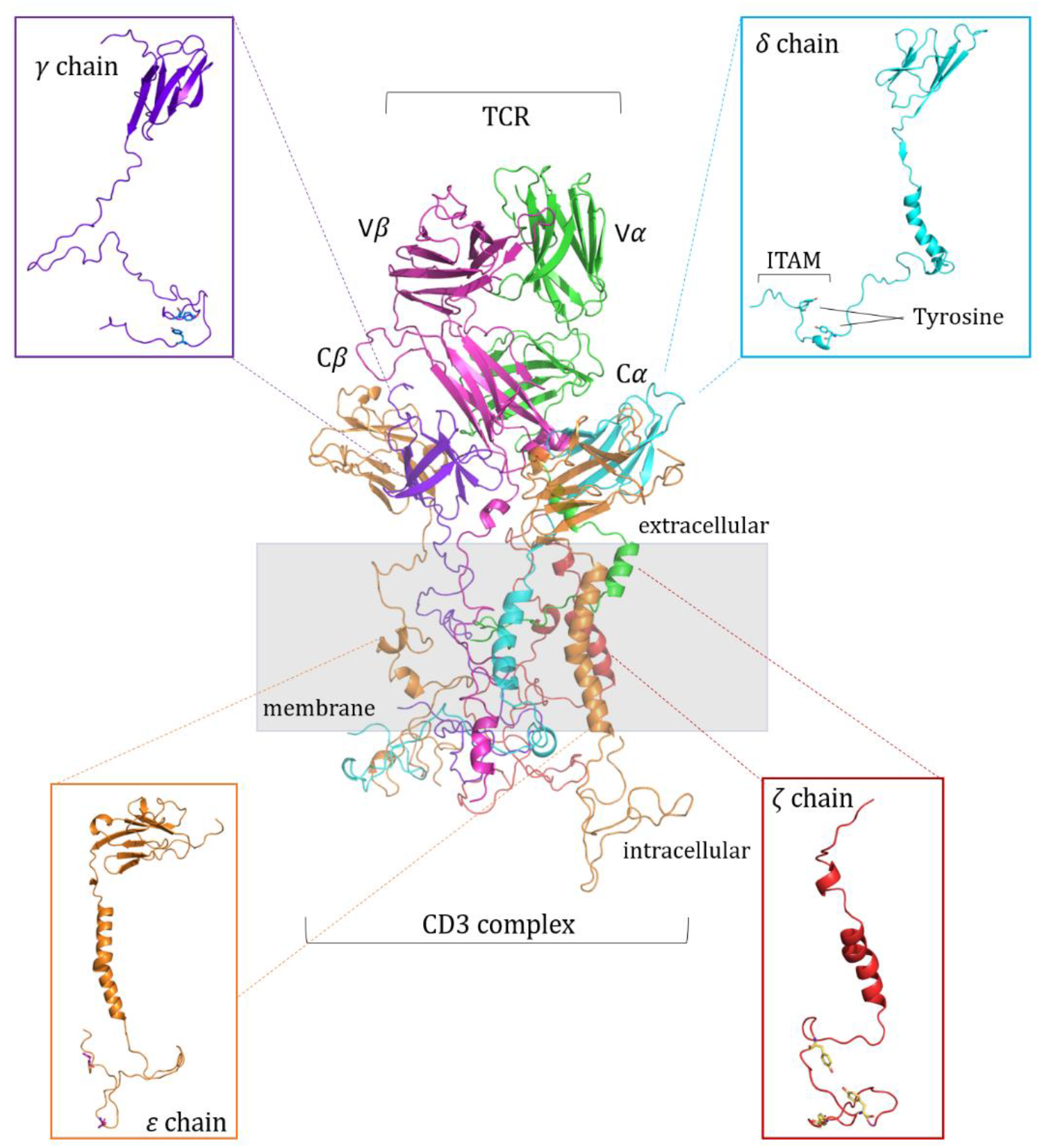
TCR:CD3-complex structure. From the cryo-EM structure [8] of the TCR:CD3-complex the missing parts were built by means of the Modeller Software. The CD3 chains interact with the terminal region of the TCR (magenta and green) through non-covalent bonds. Each chain is displayed in the box. The ITAM region, containing the tyrosines, is labelled in the *δ* chain box as reference. The TCR variable (V*α*, V*β*) and constant (C*α*, C*β*) regions are indicated.

Finally, the resulting complex was assembled and minimized in solution using the GROMACS Software version 2019.1 [12]. Then, the minimized complex was inserted manually into the membrane, precendently built by Charmm-Gui Membrane builder web software [13] with a heterogeneous lipid composition (1-palmitoyl-2-oleoyl-sn-glycero-3-phosphocholine (POPC) at 90%; phosphatidylinositol (4,5)-bisphosphate (PIP2) at 7%; 1-palmitoyl-2-oleoyl-sn-glycero-3-phospho-L-serine (POPS) at 3% [10]). A system of ∼1014000 atoms was obtained.

To compare our findings with previous experimental [14] and computational works [10] (see Table 1), we modelled the HLA-A*02:01 containing the peptide ESO9C (the tumor antigen NY-ESO157-165 fragment—SLLMWITQC [14,15]) and the mutated ESO4D (a.a. sequence SLLDWITQV [14]), interacting with the 1G4 TCR and the CD3 chains.

**Table 1.** The simulated complexes and the corresponding experimental binding affinity data [14]. The MHC class I is the HLA-A*02:01. The mutated residues are reported in red.

| Complex<br>with<br>HLA-A*0201 | Peptide<br>sequence | KD<br>(uM) | Koff (s-1) | t ½<br>(s) | Kon (M-1 s-1) |
| --- | --- | --- | --- | --- | --- |
| 1G4-ESO9C | SLLMWITQC | 14 | 0.82 | 0.84 | 57*10+3 |
| 1G4-ESO4D | SLLDWITQV | 252 | 2.59 | 0.27 | 10*10+3 |

### MD simulations

The complexes were solvated with the TIP3P water model [16] adding Na+ and Cl-ions at physiological concentration (0.15 M). The exceeding water molecules were manually removed, excluding the solvent within a range of 2 Å from lipids. An energy minimization step was performed using the steepest descent algorithm without position restraints. Following, a series of equilibration steps were carried out: (1) an NPT equilibration of 40 ps was run to allow the packing of the lipids around the protein, with an integration time step of 0.2 fs; then the NPT equilibration was extended until 2 ns. (2) an NVT equilibration of 40 ps was performed and then extended until 4 ns, increasing the time step at 1 fs. (3) A latest NPT simulation 1 us long was run with a time step of 2 fs. The Parrinello-Rahman barostat [17] and the V-rescale thermostat [18] were used with a semi-isotropic coupling with a τp = 5 ps and a τT = 0.1 ps, respectively. Considering the melting point of the lipid composition, the temperature was kept constant at 305 K. The particle mesh Ewald method [19] (cut-off of 1.2 nm) was used to treat the electrostatic interactions. A cut off of 1.2 nm was used for the van der Waals interactions. The simulations were performed using the additive for lipids CHARMM-36 force field [13,20], and the Gromacs Software version 2019.1 [12]. For each system, we performed a Molecular Dynamics simulation lasting 1us, providing several copies of the system (see below).

## Results

### Conformational Analysis

Due to the huge dimension of the complexes, two initial MD simulations of ∼600 ns were performed for each system, and one simulation of ∼700ns was provided for the unbound state.

Unexpected, in the 1G4:ESO4D:CD3 simulations, the dissociation of the HLA-A*02:peptide was observed. To investigate in detail the possibility to model such a dissociation event, we performed 12 additional MD simulations. The same event was not found for the 1G4:ESO9C:CD3 where the pMHC remains bound to the TCR for the entire simulation time (see Figure 3). Thus, no further simulations were run for this latter complex.

**Figure 3.**
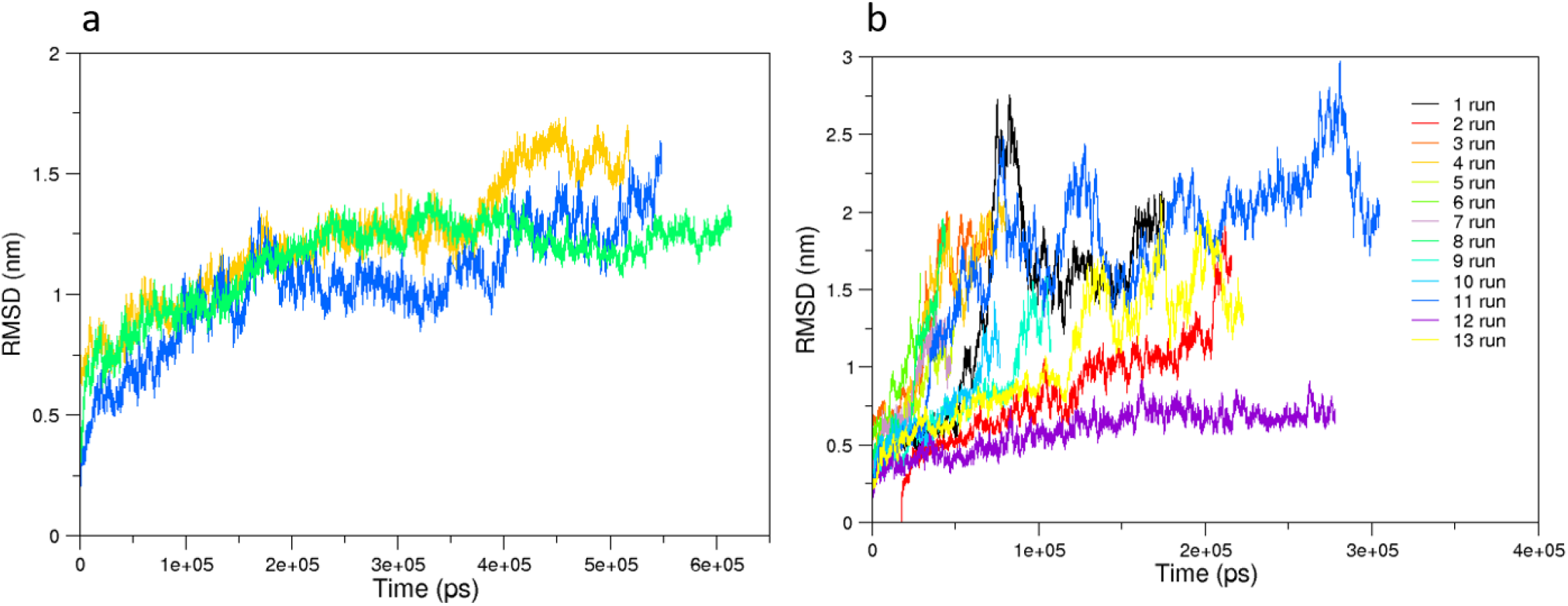
The RMSD computed on the alpha carbons of the systems. In panel a, the RMSD of the 1st run of 1G4:ESO9C:CD3 (orange), the 2nd run of 1G4:ESO9C:CD3 (blue), and of the unbound 1G4:CD3 (green). In panel b, the RMSD of the 13 trajectories of the 1G4:ESO4D:CD3 complex; in all these simulations the dissociation of the pMHC was observed, except for the run 12 (violet).

The RMSDs showed that the 1G4:ESO9C:CD3 and the unbound 1G4:CD3 reach a plateau after several ns, because of the larger dimension of the complex. Thus, the first 180 ns were removed from the analysis. To compare 1G4:ESO9C:CD3 with respect to the 1G4:ESO4D:CD3 the simulation where the dissociation is absent was considered.

The RMSF showed comparable fluctuations of the a.a. residues (Figure 4), except for i) the TCR chains, ii) the CD3*γ* and iii) the *ε*1 chains of the 1G4:ESO9C:CD3 complex (Figure 4) panel c-f). Unexpectedly, no significant differences were observed in the fluctuations of both the *ζ* chains in all the three complexes. However, 1G4:ESO9C:CD3 shown greater fluctuations of the *α* chain of the pMHC with respect 1G4:ESO4D:CD3 (Figure 4, panel i).

**Figure 4.**
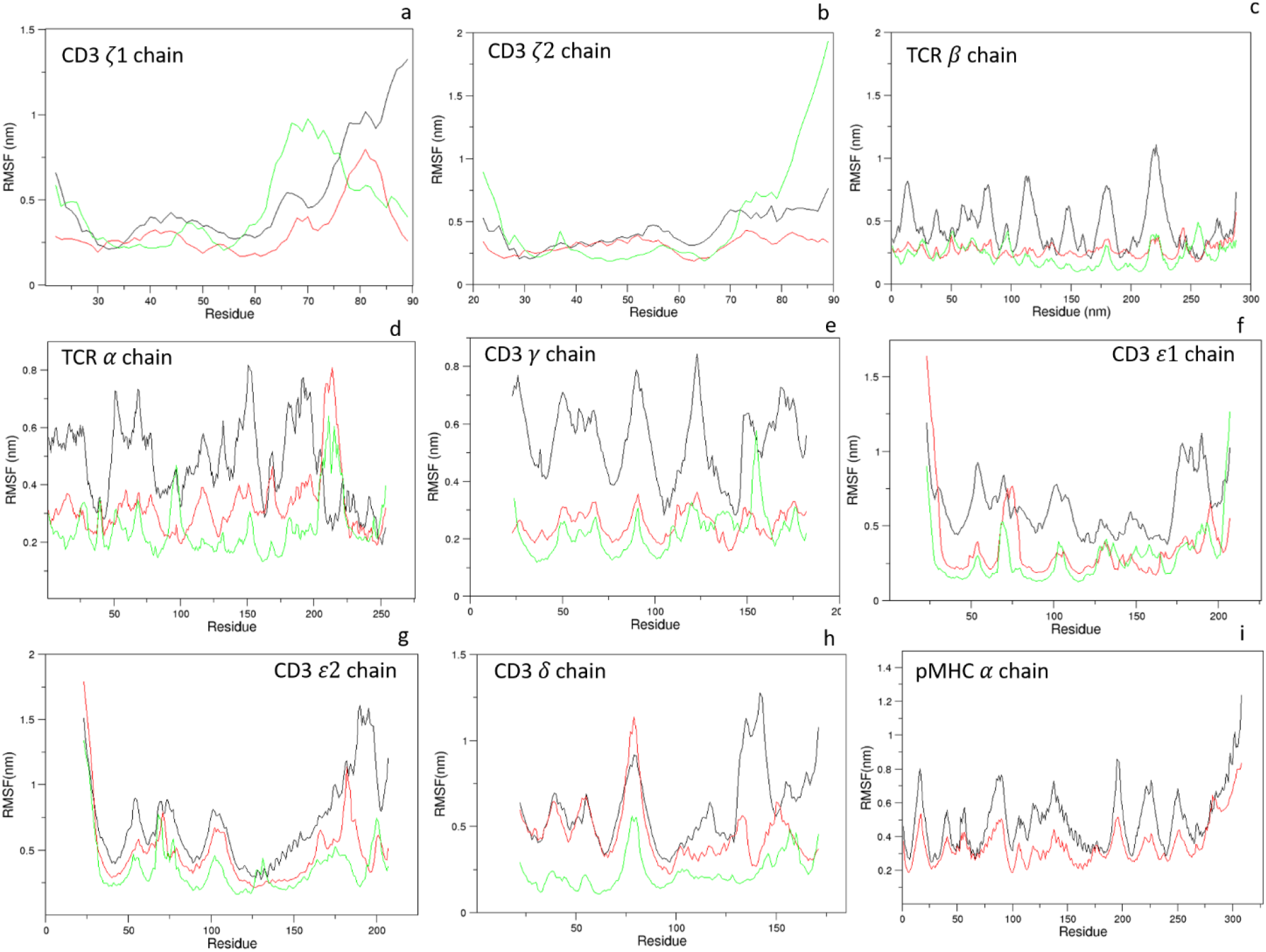
The RMSF computed on the alpha carbons of the systems. From panel a to panel h the RMSF of the CD3 chains of 1G4:ESO9C:CD3 (black), 1G4:ESO4D:CD3 (red). 1G4:CD3 unbound (green). In panel i, the RMSF of the *α* chain of the pMHC for 1G4:ESO9C:CD3 (black), and 1G4:ESO4D:CD3 (red) are shown.

### Collective motions

The overall motion of the simulated complexes was compared by means of the Essential Dynamics method. Such an analysis indicates that the first eigenvector describes about 50% of the total variance of the system. Thus, we compared the complexes with respect to the unbound 1G4:CD3, projecting on the same essential subspace the conformations explored by the MD trajectories (Figures 5-7). As expected, the projections of the alpha carbons of the TCRs show three different regions for each complex (Figure 5a), while the binding groove projections (Figure 5b) explore similar regions, in line to our previous results [10].

**Figure 5.**
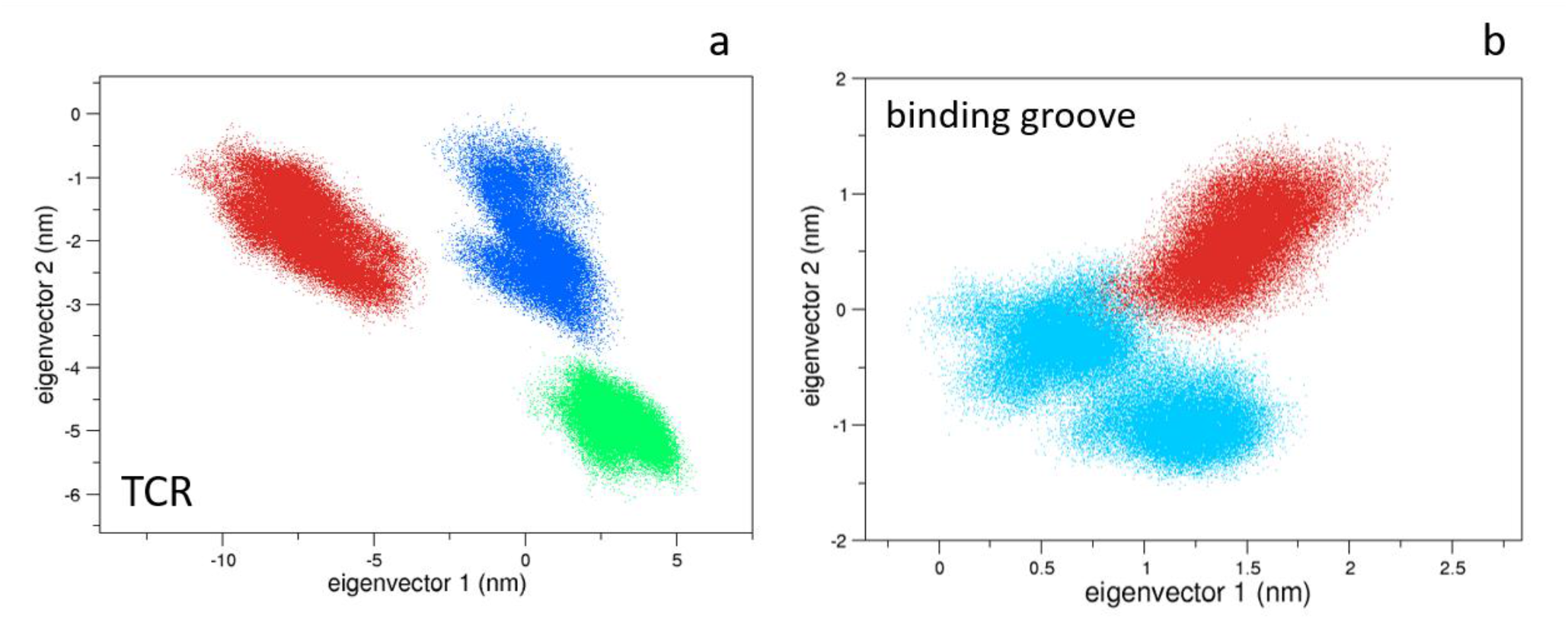
The 2D projections of TCR and Binding groove. In panel a, the projections of the TCR alpha carbons on the first 2 eigenvectors of the unbound 1G4:CD3 (green), ESO9C:TCR:CD3 (blue) and ESO4D:1G4:CD3 (red). In panel b, the projections of the HLA-A*02.01 binding groove alpha carbons of the bound states (ESO9C:1G4:CD3 in blue, ESO4D:1G4:CD3 in red).

Concerning the single TCR domain, the constant *β* region of the unbound state differs from the structures sampled by both the bound states (Figure 6d). This data, which is in line with the NMR spectroscopy outcomes [3], was not observed in our previous study [10]. This suggest that the presence of the CD3 chains modify the conformations of the TCR*β* chain. However, the conformations of the other TCR regions are almost similar, in line with our previous simulations [10]. Projecting the CD3 chains conformations on the same essential subspace, it is noticed that the unbound state explores different regions with respect to the bound states (Figure 7). Here, the first eigenvector describes about 70% of the total variance of the system, thus indicating that the CD3 chains in ESO9C and ESO4D explore similar regions of the conformational space.

**Figure 6.**
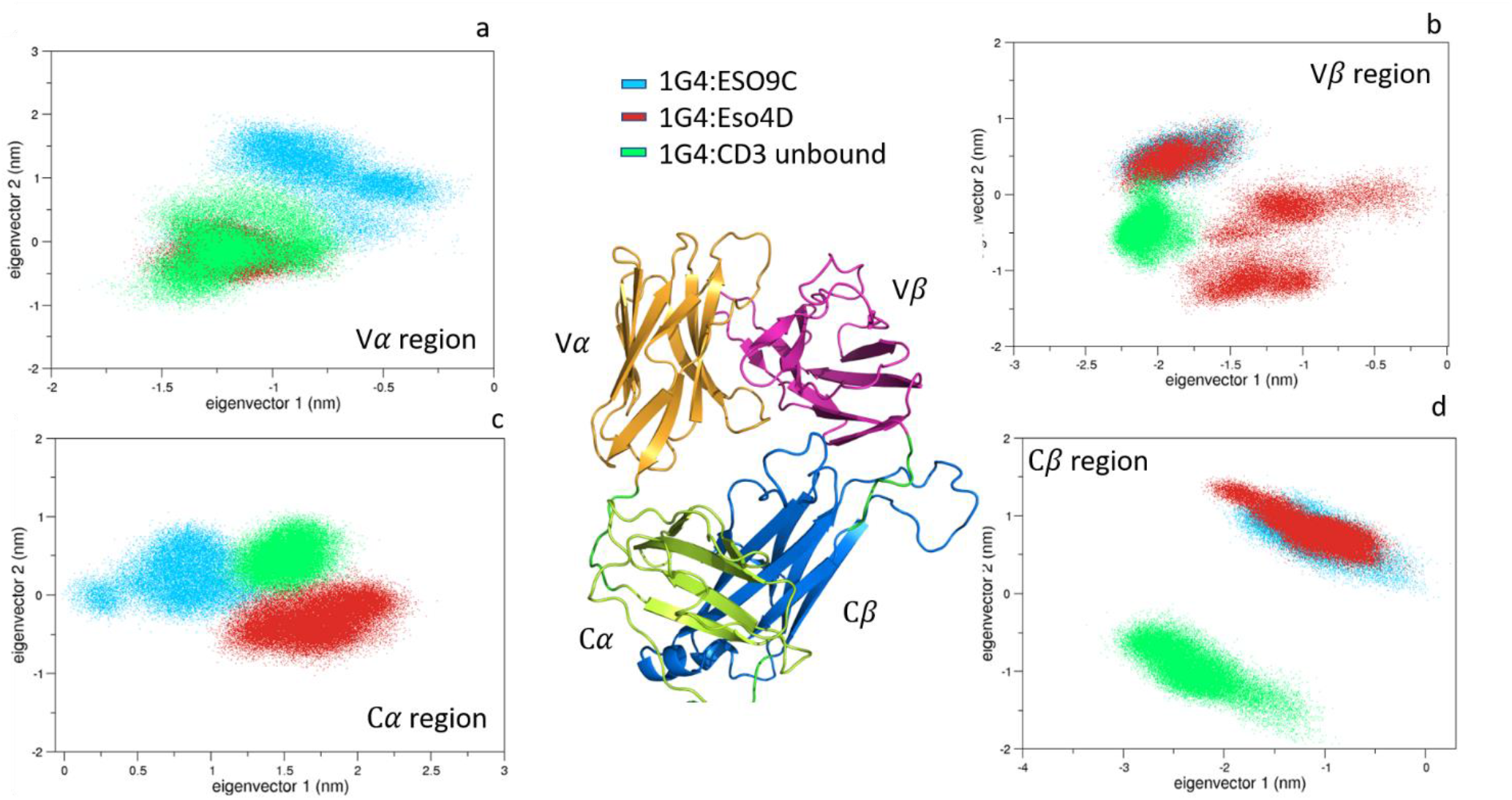
The 2D projections of the Variable and Constant regions of the TCR. The regions are labelled.

**Figure 7.**
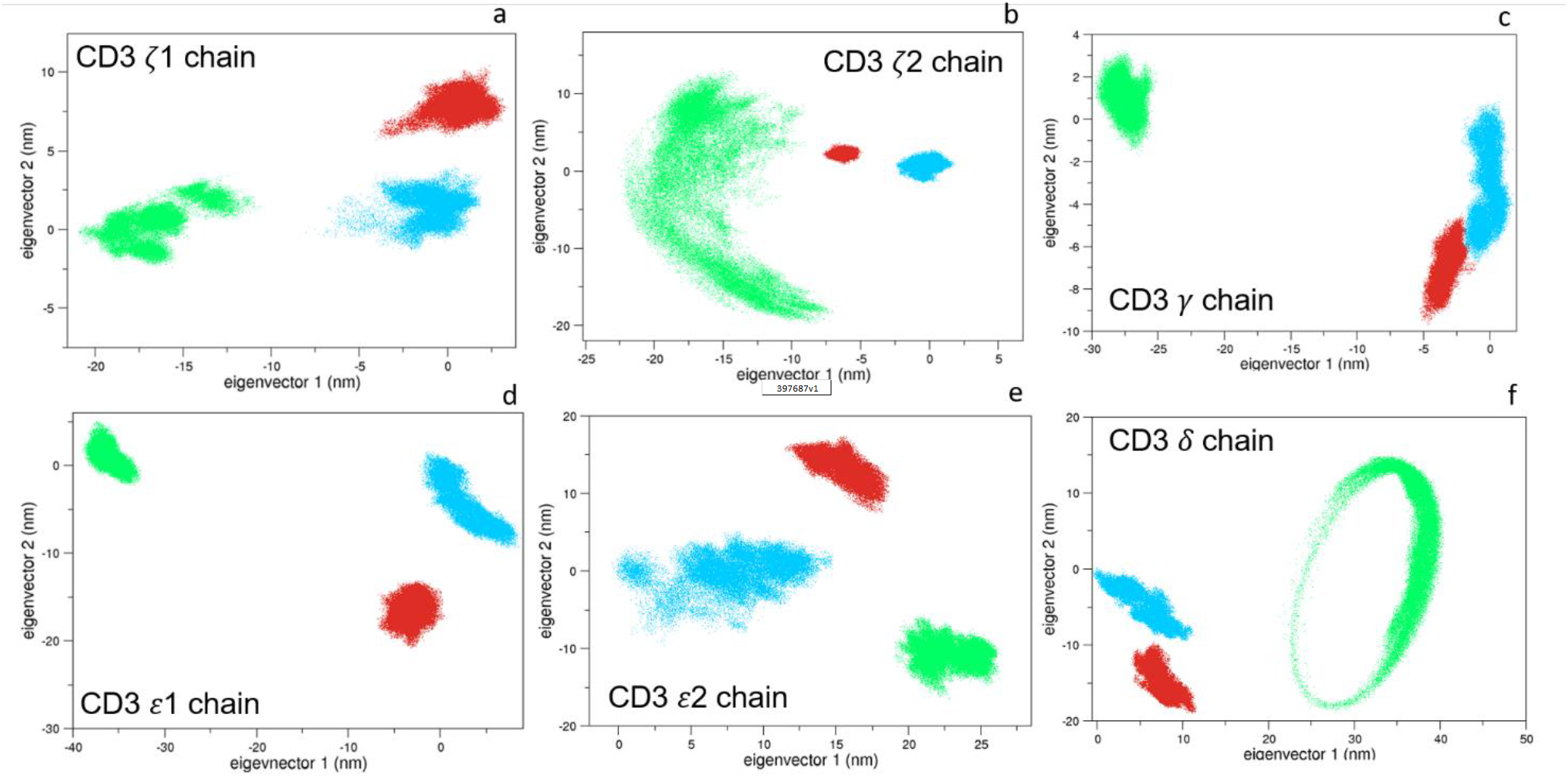
The 2D projections of the CD3 chains. The conformations of the bound states 1G4:ESO9C:CD3 (blue) and 1G4:ESO4D:CD3 (red) discriminate from the unbound 1G4:ESO4D ones (green) on the first eigenvector, which describes about the 70% of the total variance of the system.

### Dynamic Cross-Correlation (DCC) of the bound states

The DCC matrices show a greater coupling motion between the HLA-A*02.01 and the *ε*2 and the *δ* chains of the CD3 complex in 1G4:ESO4D:CD3 (Figure 8). These behaviors are missing in 1G4:ESO9C:CD3 matrix which show less or absence of correlation between the HLA and all the CD3 chains (Figure 8a).

**Figure 8.**
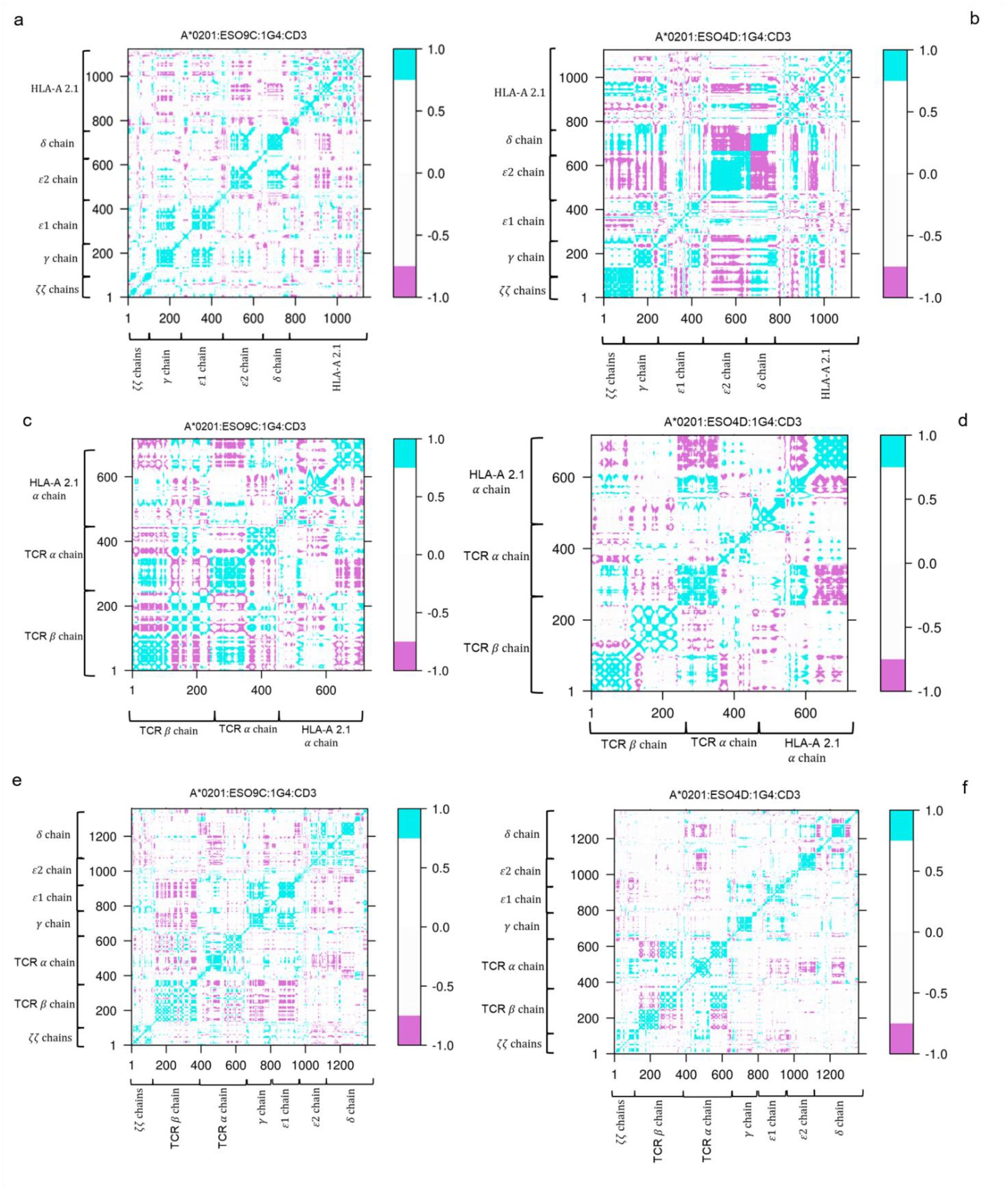
Dynamic Cross Correlation (DCC) matrices of the bound states. In panels a-b, the DCC matrices computed for the HLA-A*02.01 and the CD3 chains, for the bound states. In panels c-d, the correlations of the TCR and the CD3 chains. In panels e-f, the DCC for the TCR and the HLA-A*02.01. The cyan regions indicate the presence of a correlation; the violet regions indicate an anti-correlation; the white regions show no correlation.

Comparing the coupling between the different TCRs, it is noticed a larger correlation between the TCR*β* chain and the HLA in 1G4:ESO9C:CD3 (Figure 8c). This result is in contrast with our previous findings where the ESO4D complex shown more correlations with the TCR *β* region [10]. However, the main common coupling behavior between the TCR *α* chain is conserved in both the complexes (Figure 8c-d). Unexpectedly, no correlation was found between the TCR and the *ζζ* dimer in both the systems, but a slight coupling is reported for the TCR *α* region with all the other CD3 chains (Figure 8e-f). Interesting, a correlation between the TCR *β* and the *γ, ε*1 chains was observed for the ESO9C complex (Figure 8e). Such a correlation is missing in ESO4D, which shows an absence of coupling between the TCR-*β* and all the CD3 chains motions (Figure 8f).

### Interaction Energy of the bound states

The interaction energies between selected regions of the complexes—involving the peptide and/or the binding groove—have been analyzed (Figure 9). The distributions of such energy profiles show favorable interactions between the binding groove and the peptide in the ESO9C complex; the same interaction is shifted to higher energy value in 1G4:ESO4D:CD3. The interaction between the TCR and the peptide reports a similar profile in both the complexes, despite ESO9C -shows a bi-modal distribution of the values (Figure 9b). Finally, the interaction between the binding groove and the TCR in ESO4D displays a sharp peak of the distributions in a high energy range (Figure 9c).

**Figure 9.**
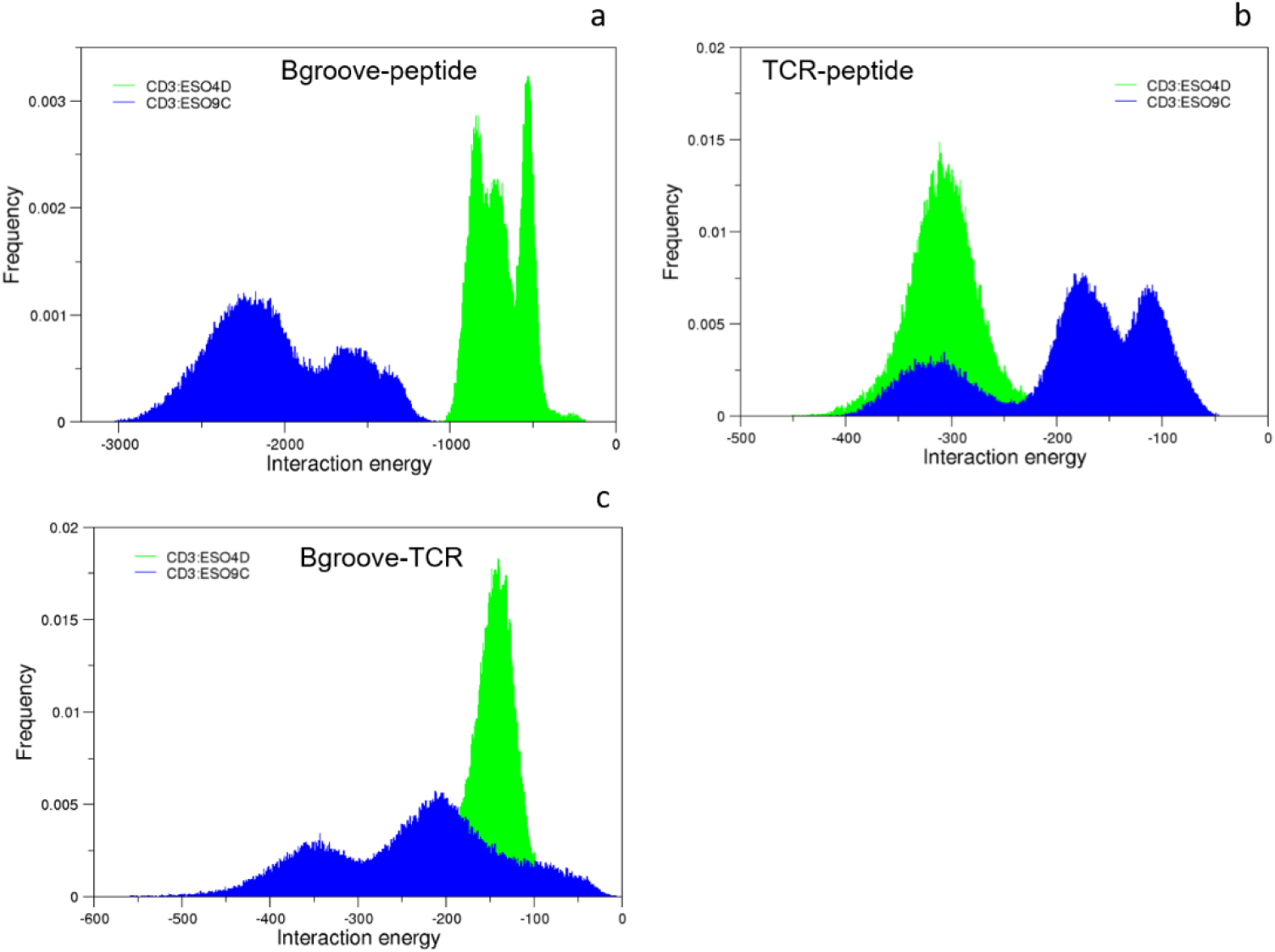
Interaction energies. The distribution of the energy profiles of the complexes is reported. Panel (a): the interaction energy between the binding groove and the peptide; panel (b): the interaction energy between the TCR and the peptide; panel (c): the interaction energy between the TCR and the binding groove.

**Figure 10.**
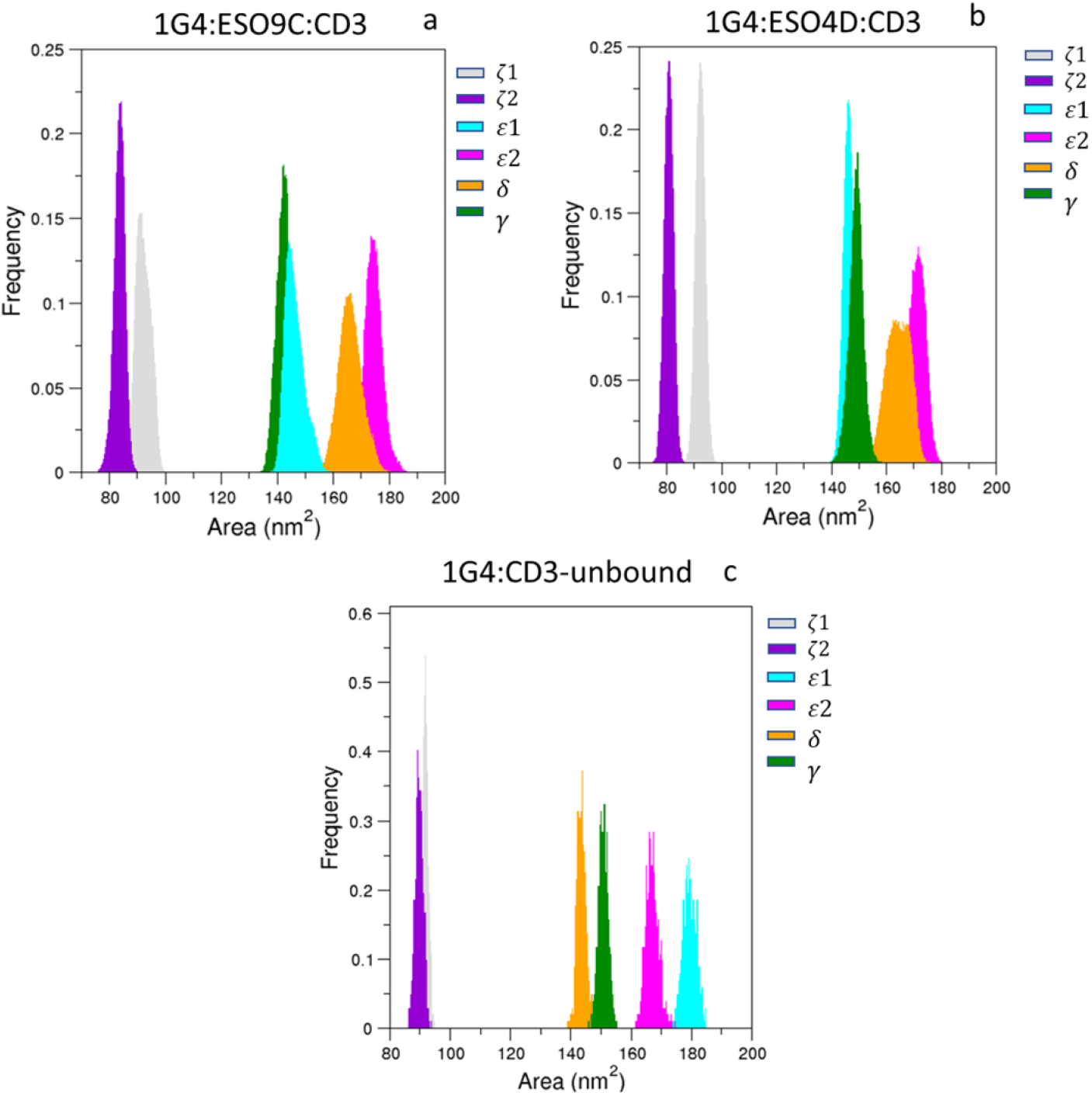
Solvent exposure of the CD3 chains.

### Solvent exposure of the CD3 chains

The solvent exposure analysis computed on the CD3 chains shows a greater exposition of the *ζ*1 and *ζ*2 chains in the 1G4:CD3 unbound. However, these results might be affected by the different lengths of the single chains. Concerning the other dimers, a larger exposure of the *δ* chains is observed in the bound states, while a lower exposition of the *ε*1 was found.

### Kinetics characterization of the 1G4:ESO4D:CD3 complex

As mentioned before, we registered several dissociation events of the complex HLA-A*0201:ESO4D from the TCR:CD3, which have been observed in 12 simulations on the 13 performed. Assuming a simple kinetic model - first order irreversible reaction - we tentatively tried to model the dissociation kinetics of such a process. The calculated kinetic constant is 0.0162 ns^−1^ (see Figure 11). It is worth mentioning that no experimental values are available for such a process.

**Figure 11.**
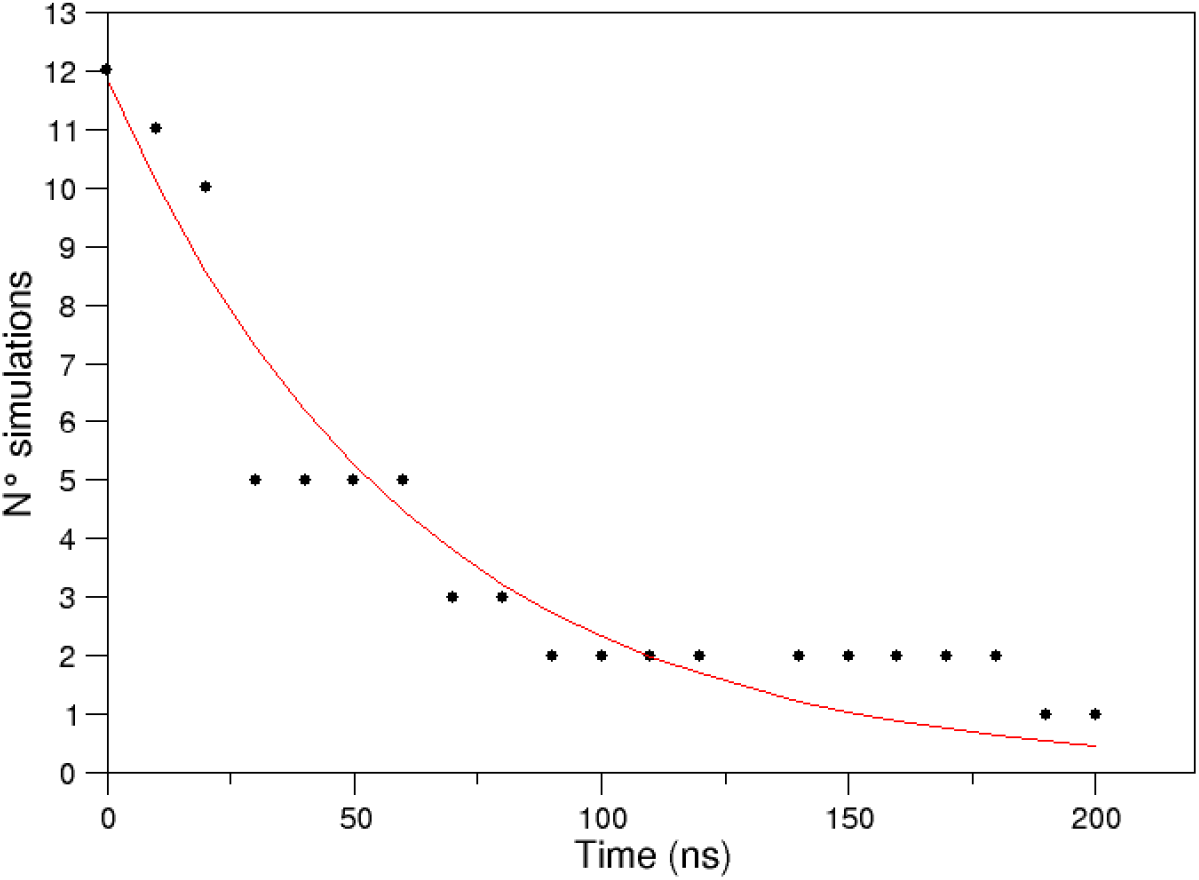
The number of the dissociation events as function of time. Black points represent the dissociation events of pMHC/TCR:CD3-complex, from the faster to the slower process. The points were fitted using the kinetic model described by formula Pop=exp(-kt). A correlation coefficient of 0.96 was obtained.

## Discussion

To consider the role of the CD3 complex in the conformational behaviour involved in the triggering process, the first computational model of the pMHC:TCR:CD3 chains was built in a lipid environment, starting from the cryo-EM structure. In this work, the same MHC class I (HLA-A*02.01) bound to two different peptides, such as ESO9C (a.a sequence SLLMWITQC) and ESO4D (a.a. sequence SLLDWITQV), were modeled binding the same TCR, as 1G4. Such systems differ from the experimentally determined kinetic constants, and from the ability of the peptide to induce the activation of the TCR, measured in terms of IFN*γ* release: in particular, ESO9C induces a greater activation of the receptor compared to ESO4D [14].

Thus, two MDs for the ESO9C complex and thirteen for the ESO4D one were carried out. In addition, a MDs of the unbound 1G4 interacting with the CD3 complex was performed, in order to compare the conformational behaviour between the bound and unbound states.

Contrary to our previous results [10], a conformational change affecting TCR C*β* region of the bound state was found. Such a change was already reported in the work of Natarajan et al. [3], by means on NMR spectroscopy. Moreover, the bound and unbound conformations of the CD3 chains explored different regions of the conformational space, while the MHC binding groove projections show a rigidity of the pocket interacting with the TCR, confirming our previous findings [2, 21].

Comparing the bound complexes, we found that the coupling motion of the TCR *α* chain with the CD3-complex is maintained in both the systems. However, only in the 1G4:ESO9C a correlation between the TCR*β* chain and the CD3*γ* and CD3*ε*1 chains was observed. No correlation was found between the motion of the TCR*β* the CD3 chains in 1G4:ESO4D, which experimentally report a minor activation of T cells. Thus, the correlation between the TCR-*β* chains and the CD3-complex could have a role in the TCR triggering.

## Acknowledgement

The authors thank Sapienza University of Rome for the financial support. We acknowledge PRACE for awarding us access to Piz Daint at CSCS, Switzerland.

